# N-Acetyl-2-Aminofluorene (AAF): *In vivo* temporal expression patterns of growth cycle-dependent macromolecular binding constants K_D[APPARENT]_ and B_MAX[APPARENT]_ are revealed by primary cultures of premalignant hepatocytes derived from a multi-cyclic hepatocarcinogenic feeding regimen

**DOI:** 10.1101/2024.09.30.615953

**Authors:** K.S. Koch, T. Moran, S. Sell, H.L. Leffert

**Affiliations:** Department of Pharmacology, School of Medicine, University of California, San Diego, La Jolla, CA 92093; Wadsworth Center, New York State Department of Health, Empire State Plaza, Albany, NY, 12201

**Keywords:** N-Acetyl-2-Aminofluorene (AAF), hepatocarcinogenesis, premalignant rat hepatocytes, primary cultures

## Abstract

Biochemical investigations of the hepatoprocarcinogen N-acetyl-2-aminofluorene (AAF) have shown that normal adult rat hepatocytes in primary culture express two sets of pharmacokinetic constants – designated Systems I and II, and Sites I and II – associated respectively with the *metabolism* (System I [high-affinity K_m[APPARENT]_ and low-velocity V_MAX[APPARENT]_] and System II [low-affinity K_m APPARENT]_ and high-velocity V_MAX[APPARENT]_]), and the m*acromolecular binding* (Site I [high-affinity *K*_D[APPARENT]_ and low capacity *B*_MAX[APPARENT]_] and Site II ([low-affinity *K*_D[APPARENT]_ and high-capacity *B*_MAX[APPARENT]_]) of AAF. Additional findings – that genomically saturating levels of AAF-DNA adducts form far below reported extracellular AAF concentrations required to block replicative and repair DNA synthesis; and, that biphasic Site I and Site II B_MAX[APPARENT]_ and K_D[APPARENT]_ expression curves varied inversely with respect to time and magnitude during hepatocyte growth – led us to wonder how macromolecular binding constants are expressed during chemical hepatocarcinogenesis. These questions were addressed by Scatchard analysis measurements through five consecutive AAF feeding cycles. Notably, cultured premalignant hepatocytes displayed reduced and elevated B_MAX[APPARENT]_ and K_D[APPARENT]_ levels, respectively, akin to the Site I and Site II expression curves observed during hepatocyte growth transitions *in vitro*. In contrast, prominent hepatocellular functions such as N-OH-AAF production, DNA replication, cell aggregation and resistance to AAF toxicity displayed different temporal trajectories.

**Impact Statement:** Striking similarities are observed between both Site I and Site II B_MAX_ and K_D_ expression curves during *in vitro* and *in vivo* premalignant growth transitions. These new findings fit earlier ones that hepatocytes growing during carcinogen exposure manifest fewer intranuclear AAF-DNA adducts. How this phenomenon leads to malignancy remains unclear.

## INTRODUCTION

Hepatocytes, the parenchymal cell population of mammalian liver, are among principal precursors to hepatocellular carcinoma (HCC) – the most frequent (80-90%) primary human liver tumor. HCC is the 2^nd^ leading cause of cancer-related death worldwide (δ > Ϙ by 3-fold); ≈ 2% of patients with distant metastases survive 5 years. HCC incidence has more than tripled over the last 4 years; this year > 800,000 people will be stricken (see 2024 websites of the American Cancer Society; National Cancer Institute; and as reviewed in Koch *et al*., 2018a,b).

Chronic infection with hepatitis B or C virus, alcoholism, smoking, and metabolic disorders like Type 2 diabetes, non-alcoholic fatty liver disease (NAFLD) and nonalcoholic steatohepatitis (NASH) are all associated etiologically with HCC. Preventive measures can ameliorate some of these conditions or reduce tumor incidence. Environmental chemicals produced by industrial and natural sources are also well-known causes of human HCC; many are *procarcinogens* which must first be metabolized to become hepatocarcinogenic. N-Acetyl-2-Aminofluorene (AAF), an occupational aromatic amine insecticide anticipated to cause human HCC (see NIOSH, 2024), is a prominent example (reviewed in Koch K.S. *et al*., 2018a).

AAF has been investigated exhaustively in animals since 1941 in attempts to explain its biochemical and biological mechanisms of hepatocarcinogenicity (Wilson, R.H., 1941; Wilson, R.H., et al., 1941); some of its principal mechanisms have been described in detail (Weisburger, J.H., et al. 1973); Becker, F.F. (ed.), 1975; Aström, A., *et al*., 1986; Barsth *et al*., 1972; Miller, E.C.,1978; Fang, D., *et al*., 2019; Weisburger, J.H., et al., 1973; Weinstein, I.B., 1978; and further reviewed in Koch *et al*., 2018a,b). AAF is initially converted enzymatically by hepatocyte-specific Cyp1A2 (a member of the cytochrome P-450 heme-thiolate mono-oxygenase superfamily) to N-hydroxy-2-acetylaminofluorene (N-OH-AAF), an N-aryl hydroxamic acid and proximal hepatocarcinogen. The reaction is selective for zone 3 liver acinar cells neighboring central veins: it requires O_2_, NADPH and reduced NADPH-hemo-protein reductase, and it occurs throughout ordered-microdomains of the cholesterol-rich endoplasmic reticulum (ER). Electrophilic metabolites of N-OH-AAF, following further modifications including deacetylation and sulfoxidation, bind covalently to hepatocyte macromolecules forming adducts with DNA, RNA, proteins and lipids. Three DNA adducts – dG-C8-AF, dG-C8-AAF and dG-N^2^-AAF – have been examined for their structural and functional properties. Some of these properties, particularly induction of error-prone DNA repair and consequent mutagenesis, are implicated in the pathophysiological changes required of chemical hepatocarcinogenesis. Yet biological evidence of which, and how, adducted DNA macromolecules cause heritable hepatocellular transformation remains unclear. For instance, several reports indicate that consensus driver mutations in three genes – *TP53*, *TERT* and *CTNNB1* – occur together at high frequency in many forms of human HCC. But the biological role(s) played by different DNA adducts in causing these and other driver mutations, or mutations in non-coding regulatory regions of hepatocyte genomes such as frame shifts or via translesion synthesis have not been definitively established (discussed in Koch *et al*., 2018b).

## MATERIALS AND METHODS

### Animals, dietary feeding regimens and feeding cycles

Male Fischer 344 rats (Simonsen Laboratories, Gilroy, CA) were used for all experiments. Customized pelleted diets were purchased from Dyets Inc. (Bethlehem, PA). Control diets (AAF^-^) were prepared without AAF; experimental diets (AAF^+^) were prepared with it (0.05% w/w). Sealed plastic bags of AAF^+^ pellets were opened inside positive pressure isolators. All rats were housed in plastic isolator cages with air filtration and access to food and water *ad libitum*.

Six-week-old rats (∼150 g) were exposed to a consecutive series of 5 feeding cycles. Each 3-week cycle consisted of 2 weeks on the AAF^+^ diet, followed by 1 week on the AAF^-^ diet (Teebor and Becker, 1971; Sell, 1978). Cycle II rats were maintained on the complete cyclic feeding regimen through cycle V and returned to AAF^-^ diets at 21 weeks of age. These rats, and separate groups continuously fed AAF^-^ diets, were also monitored for tumor formation.

Body weight and food consumption were measured as described elsewhere (Teebor and Becker, 1971; Sell, 1978; Sell, 1981). Following cycles I, III, IV and V, blood samples were drawn for assays of α1-fetoprotein (AFP [Sell, 1978]) and polypeptide hormones (Leffert *et al*., 1975). Animals were necropsied; and livers were harvested to assess gross and microscopic pathology. UCSD Institutional and Animal Care Use Committee regulations and NIH guidelines were followed for care and use of animals (Koch *et al*., 2018a).

### Primary hepatocyte culture

Single cell suspensions of adult male hepatocytes were isolated from collagenase-perfused AAF^+^ livers at the ends of dietary feeding cycles I, III, IV and V (N = 3 rats; 3 dishes/time point), or at identical times from the continuously fed group (N = 2 rats; 3 dishes/time point), and plated as described elsewhere (Leffert, H.L., *et al*., 1977, 1979). Under these plating conditions, oval and nodule cells would not have been recovered. Cycle II rats were sacrificed at death or upon tumor formation.

One million viable cells (≥ 10 µm diameter) were plated (N_0_) into uncoated 3.5 cm Falcon™ plastic tissue culture dishes (Corning/VWR) containing 2 mL arginine-free Dulbecco and Vogt’s Modified Eagle’s Medium (standard plating conditions). Culture reagents, media construction and chemicals were described previously (Leffert, H.L., *et al*., 1977, 1979). Standard plating medium was supplemented with heat-inactivated 15% v/v dialyzed fetal bovine serum (Gibco/Thermo Fisher) and 0.2 mM L-ornithine. As indicated below, some plating media were supplemented further with 10 μg insulin, hydrocortisone-succinate and inosine/mL (S^+^); or, with equivalent volumes of ornithine-supplemented serum-free medium serving as the diluent control (S^-^). Hepatocytes were cultured at 37°C in humidified 90% air-10% CO_2_ incubators. Unless otherwise stated or shown, errors of measurements using either 2 or 3 dishes per point were ±10%.

### In vitro DNsynthesis, growth and cytotoxicity assays

Attached cells were harvested from washed dishes by trypsinization and dispersed manually into single cell suspensions (viability > 95%) using 2 mL glass pipettes fitted with rubber bulbs; cell numbers were quantified using a Coulter Counter (Leffert, H.L., *et al*., 1977, 1979). Cellular attachment (cell numbers/dish) was determined 2 days post-plating. Proliferative changes were estimated by calculating the net increases in cell numbers/dish over a 7-day interval ([day 9] minus [day 2]). To reveal growth factor independence *in vitro* – a property of transformed cells, as well as a phenotype similar to reduced growth factor requirements of proliferation observed in cultured hepatocytes derived from normal rats undergoing liver regeneration post-70% hepatectomy, or from rats fed lipotrope deficient diets (Leffert, H.L., and K.S. Koch, 1978) – freshly isolated hepatocytes were split into two groups upon plating: as described above, one received supplement (S^+^) while the other received diluent (S^-^). S-phase labeling indexes (LI) were determined by autoradiography of parallel sets of cultures of similarly treated hepatocytes washed 6x with Tris-HCl buffer, pH 7.4; fixed with neutral buffered formalin; exposed to β-particle sensitive Eastman Kodak™ AR-10 stripping film; and, developed, and stained with crystal violet. DNA synthesis rates were quantified by the uptake of 1.25 μCi [^3^H]-TdR/mL (3 μM unlabeled TdR) into trichloroacetic acid (TCA)-insoluble material (cpm/106 cells/24 h). Three μM TdR was added to saturate the K_m_ of the hepatocyte TdR transport and circumvent growth factor induced changes in dTTP pools (Paul, D., *et al*., 1972).

### Cytotoxicity studies

AAF (≥ 98% purity [Sigma Aldridge]) and EtOH were dissolved in sterile serum-free medium (final concentrations of 2 x 10^-4^ M and 1% v/v, respectively) and added to cultures 1 day or 8 days post-plating. Serum-free medium containing 1% EtOH served as the diluent control. DNA synthesis rates and cell attachment measurements were made 24 h later on day 2 and day 9 post-plating.

### Isolation and quantification of soluble radiolabeled AAF metabolites

N-acetyl-9-[^14^C]-2-aminofluorene ([^14^C]-AAF; 100 dpm pmol^-1^) was purchased from New England Nuclear (Boston, MA). [^14^C]-AAF (2 x 10^-5^ M) was added to day 1 or to day 8 cultures. Cell-free culture fluids were harvested 24 h later (∼2 mL/dish); incubated with β-glucuronidase and α-amylase and extracted with acidified organic solvents (24). Radiolabeled products from processed extracts (∼65% of the counts) were assayed in parallel by thin layer chromatography (TLC) in chloroform:methanol [97:3] and benzene:acetone [4:1] as described elsewhere (24). Two TLC systems were necessary to separate 1- and 3-position ring-hydroxylated metabolites from parental AAF. Radioactivity in the samples was quantified by scintillation counting (24). The recoveries of metabolites extracted from TLC plates under both conditions were > 95%. Authentic AAF and AAF-metabolite standards were used for comparison to identify and quantify co-migrating metabolites in parallel TLC tracks (Koch *et al*., 2018a).

### Measurements of B_MAX[APPARENT]_ and K_D[APPARENT]_ in premalignant hepatocytes in primary cultures

Hepatocytes were isolated from the livers of AAF^-^ and AAF^+^ rats at the end of each feeding cycle and cultured as described above. [^14^C]-AAF was added to day 1 and to day 8 culture fluids over a wide-range of concentrations (1.25 x 10^-7^ M - 4.6 x 10^-4^ M). Twenty-four hr later (day 2 and day 9), total cell extracts were harvested from washed dishes for quantitative determinations of radiolabeled free (TCA-soluble) and covalently bound (TCA-precipitable) [^14^C]-fluorene residues. B_MAX[APPARENT]_ (B_MAX_) and K_D[APPARENT_ (K_D_) were obtained by Scatchard analyses (Koch *et al*., 2018b). Data were calculated and plotted using GraphPad™ and Prism™ software (La Jolla, CA).

### Statistical analyses

Unless noted, experimental curves were plotted without error bars, as the mean value (N) of each one their points = *XX* (results averaged from 2 or 3 rats x 3 dishes/measurement = 6 or 9) varied ± 10%. In some instances, the results were expressed as *XX* ± one standard deviation (σ) and plotted with error bars. *XX*, σ and *P* levels were calculated using Microsoft Excel™. Though hepatocyte growth transition rates decline discontinuously in aging animals (most noticeably in rats between 48-60 weeks), such physiological changes could not have influenced measurements made with AAF^-^ or AAF^+^ hepatocytes since they were isolated from younger rats at 9, 15, 18 and 21 weeks of age (cycle I, III, IV and V, respectively).

## RESULTS

### Evidence of Two Principal Constants of Macromolecular Binding Sites in Cultured Hepatocytes Isolated from Adult Male Rats Exposed to a Multi-Cycle AAF Feeding Regimen

To determine if four macromolecular binding constants revealed *in vitro* reflected similar hepatocellular expression *in vivo*, quantitative measurements of [^14^C]-AAF binding were made by Scatchard analyses in cultured hepatocytes spanning growth cycles of primary cultures of hepatocytes isolated from rats fed either normal chow diets (AAF^-^) or 1 to 5 cycles of AAF-supplemented hepatocarcinogenic diets (AAF^+^). Measurements were obtained with cycle I, III, IV and V hepatocytes on day 2 and day 9 post-plating; culture media in each group contained growth factor supplement (S^+^) or diluent (S^-^). Figure 1 shows that macromolecular binding sites defined by low-capacity B_MAX[APPARENT]_ and high-affinity K_D_ (Site I) and by high-capacity B_MAX[APPARENT]_ and low-affinity K_D_ (Site II) were observed.

**Figure 1.**
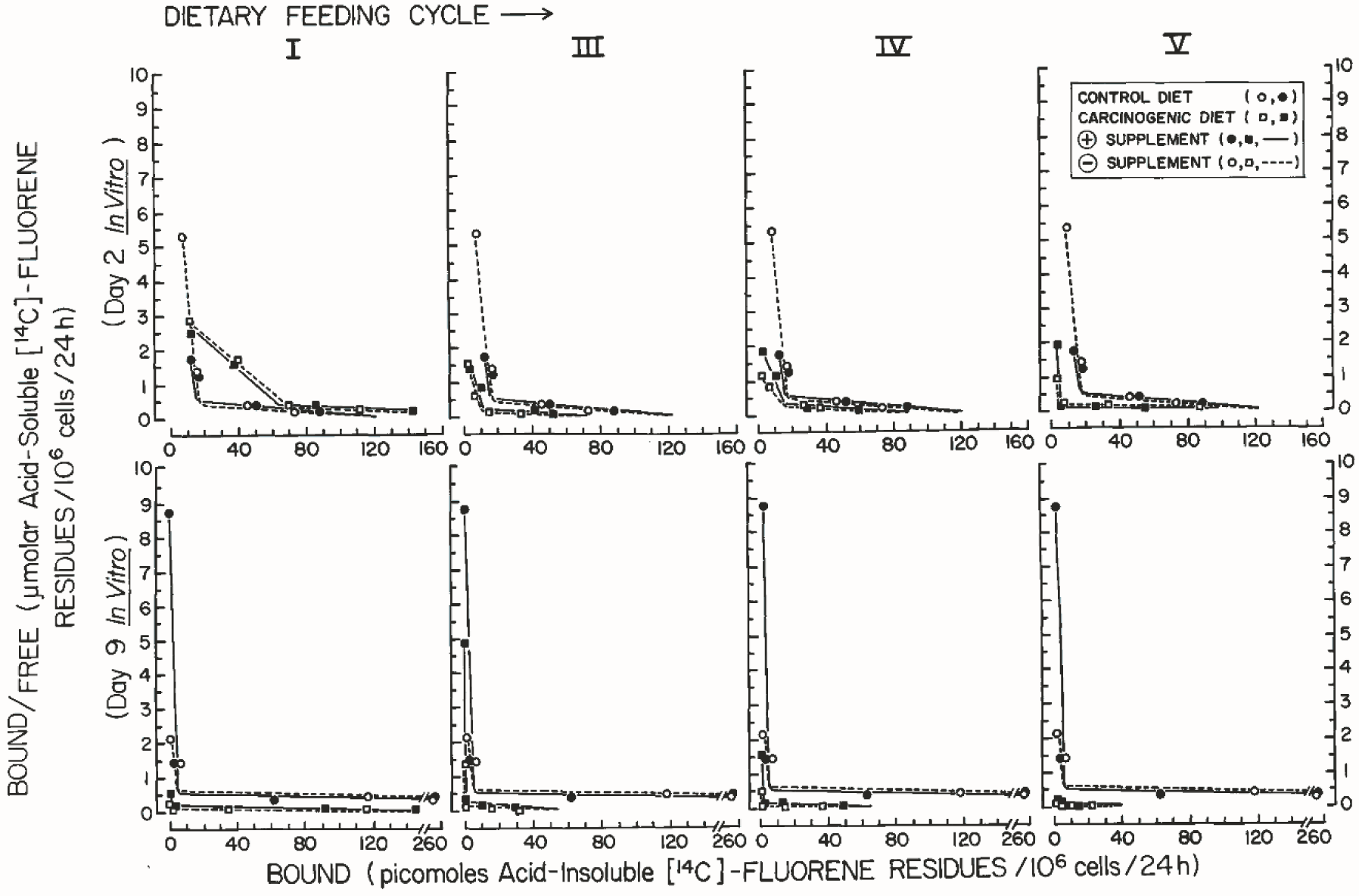
Quantitative binding of [^14^C]-AAF reveals two constants of AAF macromolecular binding sites in primary cultures of hepatocytes derived from rats exposed to a multi-cycle AAF feeding regimen. Primary cultures of hepatocytes from cycle I, III, IV and V rats – fed control AAF^-^ (o, ●) or carcinogenic AAF^+^ diets (□, ▪) – were plated as described in Material and Methods. Symbols (inset box [far upper right panel]) also denote media which were supplemented with 50 ng insulin, inosine and hydrocortisone-succinate/mL ( ) or with diluent (o - - - - □). *In vitro* assays of bound and free [^14^C]-fluorene residues were performed 2 days (top panels) and 9 days post-plating (bottom panels) as described in Material and Methods. The resulting curves were used to calculate K_D_ and B_MAX_ (Koch *et al*., 2018b). For all cycles, the results from control cultures (AAF-S- [o, o] or AAF-S+ [●,●]) were combined and averaged (N = [2 rats] x [4 cycles] x [3 dishes/point]) = 24 samples); the resulting composite curve was plotted alongside experimental curves (AAF+S- [□- - - □] or AAF+S+ [▪, ▪]).

### Distinct Temporal Patterns of AAF Binding to Macromolecular Sites I and II are Revealed in Premalignant Hepatocytes Exposed to a Multi-Cycle AAF Feeding Regimen

Site I and II B_MAX[APPARENT]_ and K_D[APPARENT]_ data from Figure 1 were replotted as time-courses to visualize premalignant changes representing all four constants during the 15-week regimen. The results are shown in Figure 2 for cycle I, III, IV and V hepatocytes: B_MAX[APPARENT]_, Site I and Site II (low- and high-capacity [panel A, left and right sides, respectively]); K_D_, Site I and Site II (high- and low-affinity [panel B, left and right sides, respectively]).

**Figure 2.**
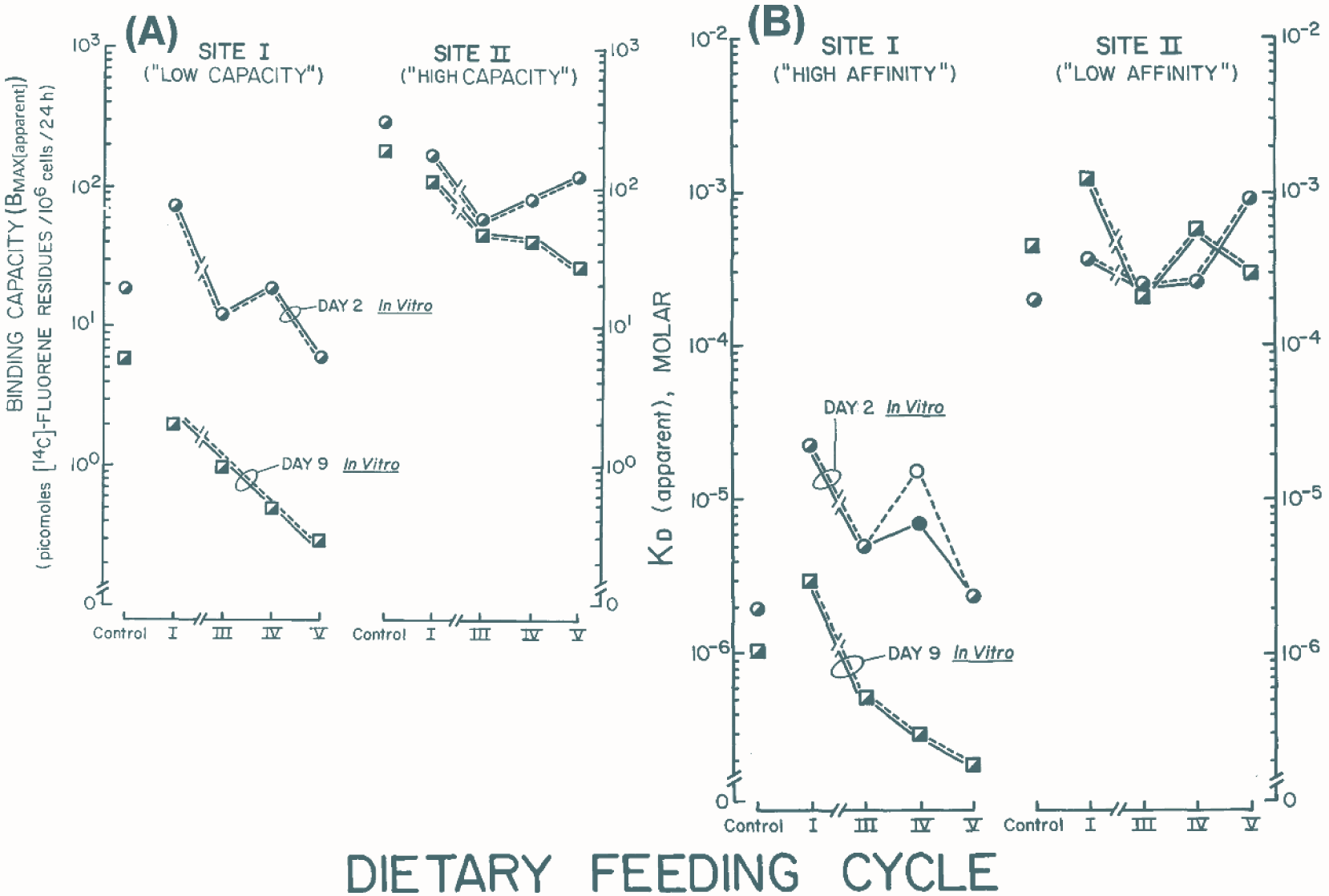
Time course plots reveal premalignant temporal expression patterns of [^14^C]-AAF binding to macromolecular Sites I and II. Hepatocytes from cycles I, III, IV and V were plated with (S^+^) or without supplement (S^-^) and assayed 2 days (o, ●) and 9 days (□, ▪) post-plating as described in Material and Methods. Data from Figure 1 were used to calculate two pharmacokinetic constants: B_MAX[APPARENT],_ binding capacity (picomoles [^14^C]-fluorene residues/10^6^ cells/24 h); and, K_D[APPARENT]_, (Molar). A, Site I B_MAX[APPARENT]_, low-capacity (left side); and Site II B_MAX[APPARENT]_, high-capacity (right side). B, Site I K_D_, high-affinity (left side); and, Site II K_D_, low-affinity (right side). For each one of the 8 curves, the results from control cultures (AAF^-^S^-^ or AAF^-^S^+^ [day 2 or day 9]) were combined, averaged (N = [2 rats] x [4 cycles] x [3 dishes/point] = 24 samples) and plotted as single unconnected points to the left of each curve. All points show split white-and-black shading because the results of S^-^ and S^+^ groups were statistically identical.

Compared to S^-^ controls, but for one anomalous cycle IV point involving a day 2 measurement of the AAF^-^ Site II high-affinity K_D_ (where S^-^ > S^+^), none of the curves were affected by growth factor supplementation of the culture media. Also, regarding AAF^-^ control culture measurements made on day 2 and day 9 post-plating, with the exception of the Site II low-affinity K_D_ which fell ∼ 2-fold (2 x 10^-4^ M to 4 x 10^-4^ M), the trends observed for AAF^-^ control cultures between day 2 and day 9 post-plating (Site I low-capacity B_MAX[APPARENT]_, 20 to 6; Site II high-capacity B_MAX[APPARENT]_, 300 to 200; Site I high-affinity K_D_ (2 to 1 x 10^-6^ M) were similar to those reported for primary cultures of normal hepatocytes (Koch *et al*., 2018b).

Distinct temporal patterns were observed among the four sets of curves. Initially, compared to AAF^-^ controls, the constants of several cycle I hepatocytes were elevated. Three sets declined: Site I low-capacity B_MAX_ (panel A, left), which followed a normal hepatocyte growth cycle trend (Koch *et al*., 2018b); Site II high-capacity B_MAX_ (panel A, right); and Site I high-affinity K_D_ (panel B, left), which also followed a similar growth cycle trend (Koch *et al*., 2018b). The fourth set – Site II low-affinity K_D_ – remained relatively invariant with time (panel B, right). Day 2 measurements quantitatively exceeded day 9 levels in the 1^st^ three sets, but not in the fourth. Thus, from day 2 to day 9, control Site I levels of low-capacity B_MAX_ and high-affinity K_D_ fell from 19 to 6 picomoles/10^6^ cells/24 h (3.2-fold) and 2 x 10^-6^ to 1 x 10^-6^ M (2-fold), as shown in Figure 2A and Figure 2B, respectively. Concomitantly, Site II levels of high-capacity B_MAX_ levels fell from 300 to 195 picomoles/10^6^ cells/24 h (1.54-fold), while low-affinity K_D_ levels rose from 2 x 10^-4^ to 4.5 x 10^-4^ M (2.25-fold).

Specifically, compared to AAF^-^ cultures, from day 2 to day 9, the Site I levels of the BMAX rose from 20 to 80 picomoles/10^6^ cells/24 h (Figure 2A), whereas the K_D_ of cycles I-V AAF^+^ cultures fell from 2.2 x 10^-5^ M to 3 x 10^-6^ M (Figure 2B). Subsequently, from day 2 to day 9, the Site I and Site II levels of the B_MAX_ fell significantly, as a function of feeding cycle number through cycle V, from 80 to 6, and 20 to 0.3 picomoles/10^6^ cells/24 h), respectively, and from 300 to 100, and 150 to 25 picomoles/10^6^ cells/24 h, respectively (Figure 2A). Concomitant levels of the K_D_ from day 2 to day 9 did not parallel these trends through feeding cycle V (K_D_ fell 3.2-fold (19 to 6 picomoles/10^6^ cells/24 h) and 2-fold (2 x 10^-6^ to 1 x 10^-6^ M), respectively, as shown in Figure 2A and Figure 2B, respectively. Concomitantly, Site II levels of high-capacity BMAX levels fell 1.54-fold (300 to 195 picomoles/10^6^ cells/24 h), in contrast to prior findings which showed a 2.1-fold increase [Koch *et al*., 2018b]), while low-affinity KD levels rose 2.25-fold (2 x 10^-4^ to 4.5 x 10^-4^ M). Unlike elevations observed previously in control low-affinity KD levels of Site II (Koch *et al*., 2018b), no definitive trends occurred in this parameter, which fluctuated among AAF^+^ groups between 2 x 10^-4^ M - 1 x 10^-3^ M in both day 2 and day 9 cultures during the 15-week regimen covering AAF^+^ cycles I, III, IV and V.

None of these patterns could have been affected by the molar concentrations of plasma AAF over the 15-week regimen (feeding cycles I-V). Using the following information (MW AAF = 223; 0.05% w/w dietary AAF; body weight [BW]; diet consumption of g/rat/week; and, blood volume [BV]/rat (mL) = 0.06 X BW + 0.77), molar concentrations were calculated step-wise (g diet/consumed /rat/cycle → BW/BV per rat/cycle → g/AAF/rat/cycle → g/AAF/L/rat/cycle) to give the following levels for cycle I (4.78 x 10^-3^ M = 100%), cycle II (3.73 x 10^-3^ M), cycle III (3.10 x 10^-3^ M), cycle IV (3.56 x 10^-3^ M) and cycle V (3.56 x 10^-3^ M).

### AAF Metabolism and Metabolite Secretion in Cultures of Primary Hepatocytes Isolated from Adult Male Rats Exposed to a Multi-Cycle AAF Feeding Regimen

AAF metabolism (2 x 10^-5^ M) was stimulated 2-3-fold in supplemented control cultures 2 and 9 days post-plating (Figure 3).

**Figure 3.**
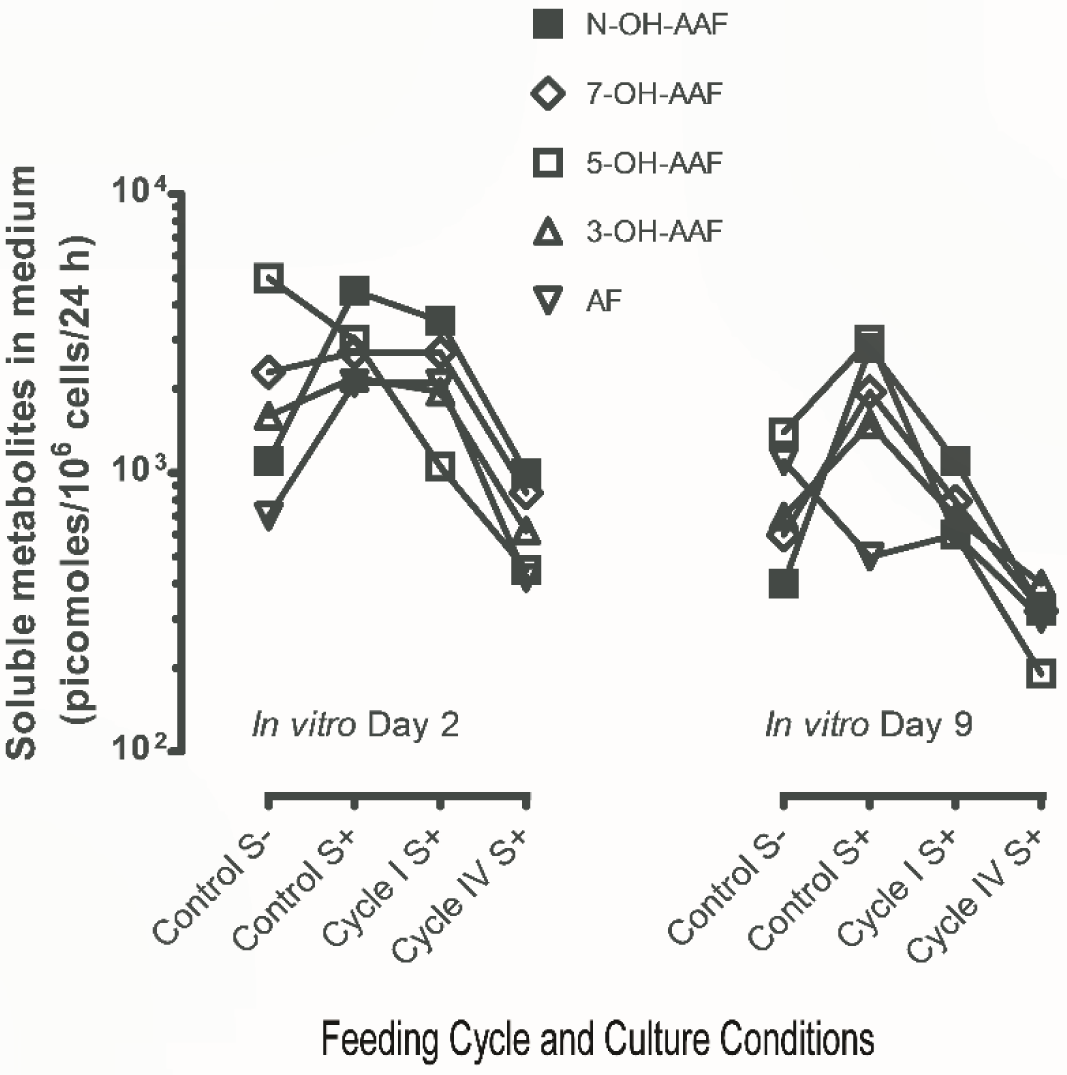
Metabolism of [^14^C]-AAF in primary cultures of hepatocytes derived from rats exposed to a multi-cycle AAF feeding regimen. AAF^-^ and AAF^+^ hepatocytes were isolated from cycle I and cycle IV rats and plated without (S^-^) or with supplement (S^+^). On day 1 and day 8 post-plating, the cultures were incubated 24 h with 2 x 10^-5^ M [^14^C]-AAF (40 nmols/4.3 x 10^6^ cpm/2 mL/culture) and 2 days (left panel) and 9 days (right panel) post-plating soluble metabolites released into the culture media were quantified (picomoles/10^6^ cells/24 h; Koch *et al*., 2018a): ▪, N-OH-AAF; ◊, ring-hydroxylated 7-OH-AAF; □, 5-OH-AAF and Δ, 3-OH-AAF; and, □, AF [de-acetylated 2-aminofluorene]). The AAF consumed/culture (d 2 ≈ d 9): AAF^-^S^-^ (50%) and AAF^-^ S^+^(52%) controls: 52%; AAF^+^S^+^ cycle I (36%) and AAF^+^S^+^ cycle IV (15%). The results from control cultures (AAF^-^S^-^ or AAF^-^S^+^) were combined and averaged (N = [2 rats] x [2 cycles] x 3 dishes/point] = 12 samples). Experimental results from AAF^+^S^+^ cycles I and IV were averages of N = [3 rats] x [2 cycles] x 3 dishes/point] = 18 samples.

Compared to unsupplemented cultures, control AAF^-^S^+^ hepatocytes generally secreted higher levels of measurable metabolites (except for the higher levels of 5-OH-AAF on day 2, and the lower levels of AF [aminofluorene] on day 9).

However, cultured hepatocytes from AAF^+^S^+^ rats lost their capacity to metabolize AAF in proportion to the time of dietary exposure to AAF. This was observed in two ways. First, over 24 h periods between 1-2- or 8-9 days post-plating, AAF^-^S^-/+^ cultures treated with AAF metabolized 50-52% of the extracellular procarcinogen; while cultured AAF^+^S^+^ hepatocytes from dietary cycles I and IV metabolized 35% and 15%, respectively (see legend to Figure 3). Steep declines in the rates of secretion of AAF metabolites were observed in AAF^+^S^+^ cultures. These results were documented for activated N-hydroxylated-AAF (▪, N-OH-AAF); ring-hydroxylated (◊, 7-OH-AAF, □, 5-OH-AAF and Δ, 3-OH-AAF); and de-acetylated products (□, AF), as shown in Figure 3. In day 9 cultures, the rates of AF secretion were much slower than the rates of the other 4 metabolites – all of which appeared to decline with 1^st^ order kinetics. The AAF consumed/culture (d2 ≈ d9): Control: 52% (2.16 x 10^6^ cpm); cycle I (∼36% [1.5 x 10^6^ cpm]) and cycle IV (15% [6.4 x 10^5^ cpm]). When 2 x 10^-5^ M [^3^H]-AAF was added to day 3 cultures, 54% of the radiolabeled extracellular AAF disappeared from the culture medium within 24 h. The rate of disappearance followed zero-order kinetics (2.3 nmol AAF/106 cells/h ∼12 µg AAF/10^6^ cells/day).

### AAF metabolism and metabolite secretion: Relationships to changes in Site I B_MAX_ and Site II K_D_

The day 2 bell-shaped N-OH-AAF metabolite secretion curve (Figure 3) followed bell-shaped Site I B_MAX_ curve. In contrast, although the Day 9 N-OH-AAF metabolite secretion curve was also bell-shaped, the B_MAX_ curve fell continuously more than 20-fold, with 1^st^ order kinetics (Figure 2A).

### Proliferative Properties of Cultures of Primary Hepatocytes Isolated from Adult Male Rats Exposed to a Multi-Cycle AAF Feeding Regimen

Figure 4 shows growth patterns of AAF^-^ and AAF^+^ hepatocytes isolated from cycle I, III, IV and V rats (the protocol’s cyclic feeding and cell culture regimens are diagrammed across the figure’s top margin, left-to-right).

**Figure 4.**
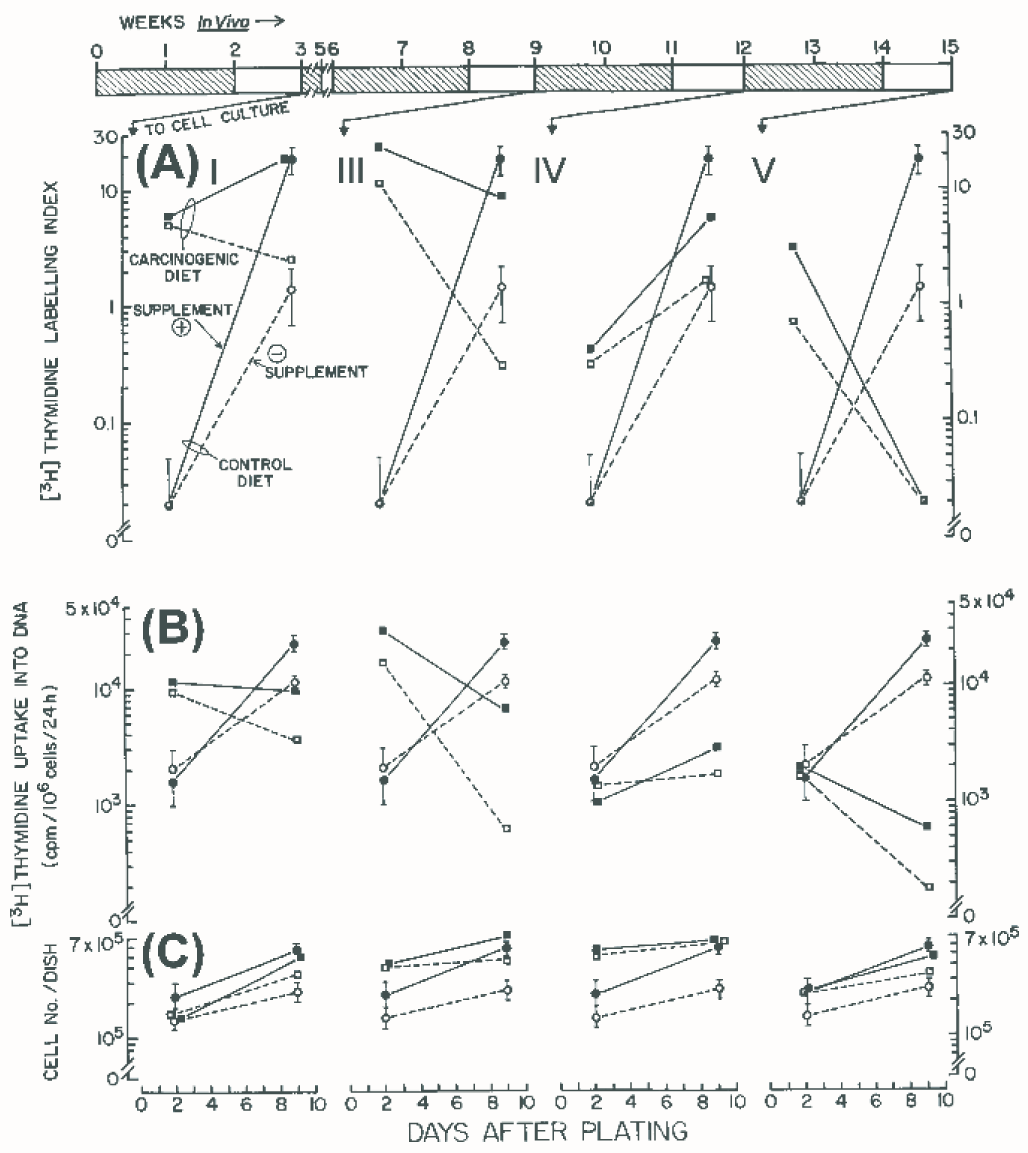
Proliferative properties of hepatocytes in primaultures derived from rats exposed to a multi-cycle AAF feeding regimen. Hepatocytes from AAF^-^ (●, ○) and AAF^+^ (▪, □) animals were isolated at the end of each indicated 3-week feeding cycle (plus AAF for 2 weeks [ ]; minus AAF for one week [ ]). This is denoted by 4 downwardly directed arrows (‘TO CELL CULTURE’ [left to right]) from the horizontal timeline bar at the top (0-15 weeks *in vivo* →). Cells from cycles I, III, IV and V were plated into standard culture media without (○- - -□) or with (●- - -▪) supplement. As shown on the y-axes, ( A) [^3^H]-TdR LI (labeling indexes [% of hepatocytes in S-phase]), (B) [^3^H]-TdR uptake into DNA, and (C) cell numbers/dish, were measured 2 and 9 days post-plating (x-axis). For all cycles, the results from control (AAF^-^S^-^ [o- - - - o] or AAF^-^S^+^ [● - -●]) cultures were combined and averaged (N = [2 rats] x [4 cycles] x [3 dishes/point]) = 24 ± σ (standard error bars) samples); the resulting composite curve was plotted alongside each one of the individual cycle I, III, IV and V experimental curves (AAF^+^S^-^ [□- - - - □] or AAF^+^S^+^ [▪- - -▪]), where N = [3 rats] x [4 cycles] x [3 dishes/point]) = 36 ± σ.

Four constants were quantified with respect to time: [^3^H]-LI (A); DNA synthesis (B); and, cell attachment (day 2) and proliferative change (C). [^3^H]-LI rates depended upon culture age, supplement and feeding cycle number. Two days post-plating, [^3^H]-LI rates in AAF^+^S^+^ cultures from cycle I, III, IV and V were 300-, 1200-, 20- and 160-fold higher (6, 24, 0.4 and 3.2%, respectively) than AAF^-^S^+^ cultures (0.02 ± 0.05%). LI rates also peaked and declined in AAF^+^S^-^cultures, but at levels 250-, 600-, 10- and 35-fold higher (5, 12, 0.2, 0.7%, respectively) than AAF^-^S^-^ cultures (0.02%; A). LI rates trended differently 9 days post-plating. AAF^+^S^+^ LI rates fell in cycle I, III, IV and V cultures 900-, 450-, 350- and 1-fold (18, 9, 7 and 0.02%, respectively), as AAF^-^S^+^ LI rates rose 900-fold to 18%. Likewise, AAF^+^S^-^ LI rates fell from 110-, 15-, 60- and 1-fold (2.2, 0.3, 1.2 and 0.02%, respectively), as AAF^-^S^-^ LI rates rose 60-fold to 1.2% (A).

As expected, DNA synthesis rates (B) were proportional to LI rates (A) in 2- and 9-day old cultures in all four *in vitro* treatment groups across all feeding cycles. Attachment and its dependence upon supplement were measured 2 days post-plating across all feeding cycles: AAF^+^ cells were unresponsive, whereas AAF^-^ cells were stimulated ∼ 2-fold (2.2 x 10^5^ [S^+^] vs. 1.3 x 10^5^ [S^-^] cells (C). Notably, AAF^+^ attachment exceeded AAF^-^ attachment by 2-3.5- and 2.6-4.4-fold during cycles III and IV (4.5 x 10^5^ and 5.7 x 10^5^ [S^-/+^], respectively); these differences were not observed during cycles I and IV, when AAF^+^ cell attachment ranged between 1.3-2.2 x 10^5^ [S^-/+^]. Proliferative change, estimated by calculating net increases in cell numbers/dish over a 7-day interval *in vitro* ([day 9] minus [day 2]), was averaged for each treatment group across all feeding cycles (C). Supplement stimulated proliferative change within AAF^-^ and within AAF^+^ groups: 4.3-fold (AAF^-^S^+^ > AAF^-^S^-^ cells [3.0 x 10^5^ vs. 0.7 x 10^5^ cells, *P* < 0.005]) and 2.7-fold (AAF^+^S^+^ > AAF^+^S^-^ [2.25 ± 1.05 (σ) x 10^5^ vs. 0.82 ± 0.48 (σ) x 10^5^ cells, *P* < 0.05]), respectively. Although supplement-responsive, differences in proliferative change were not observed between AAF^-^ and AAF^+^ groups: AAF^-^S^+^ ∼ AAF^+^S^+^ (3.0 x 10^5^ vs. 2.25 ± 1.05 (σ) x 10^5^ cells, NS) and AAF^-^S^-^ ∼ AAF^+^S^-^ (0.7 x 10^5^ vs. 0.82 ± 0.48 (σ) x 10^5^, NS), respectively.

### Aggregation Properties of Cultures of Primary Hepatocytes Isolated from Adult Male Rats Exposed to a Multi-Cycle AAF Feeding Regimen

Normal hepatocytes in primary culture form multicellular monolayer aggregates. As observed by phase microscopy, these aggregates consist of three or more hepatocytes that make membranous contacts within a defined cross-sectional area of the surface of the dish. Aggregates develop within 24 h post-plating from mono- and bi-nucleated hepatocytes that flatten upon the dish. During the ensuing growth cycle, hepatocytes morphologically undergo a reversible epithelial □ mesenchymal-like transition (Supplementary Figure 1). This transition affects aggregation (which diminishes during the mesenchymal phase) and is accompanied by reversible levels in the expression of differentiated functions and bile canaliculi. Two days post-plating, 74% of control AAF^-^S^-^ hepatocytes were situated in aggregates (6.5 ± 0.5 [σ] cells/40,000 µm^2^) [see Figure 5A]). The remaining cells consisted of non-aggregated hepatocytes (21%) and non-parenchymal cells (5%) that survived by cross-feeding. Four days post-plating, heightened spreading was observed (Supplementary Figure 1); aggregation fell significantly (3.2 ± 0.8 cells/40,000 µm^2^; *P* < 0.0004) as cells proliferated during log phase. Hepatocytes entered stationary phase 9 days post-plating, following a 200-300% increase in cell numbers/dish (Figure 4); nine days later (18 days post-plating) aggregation rose significantly (9.3 ± 0.4 cells/40,000 µm^2^; *P* < 0.0001) by 43% and 66% compared to cultures 2- and 4-days, respectively, post-plating (Supplementary Figure 1).

**Figure 5.**
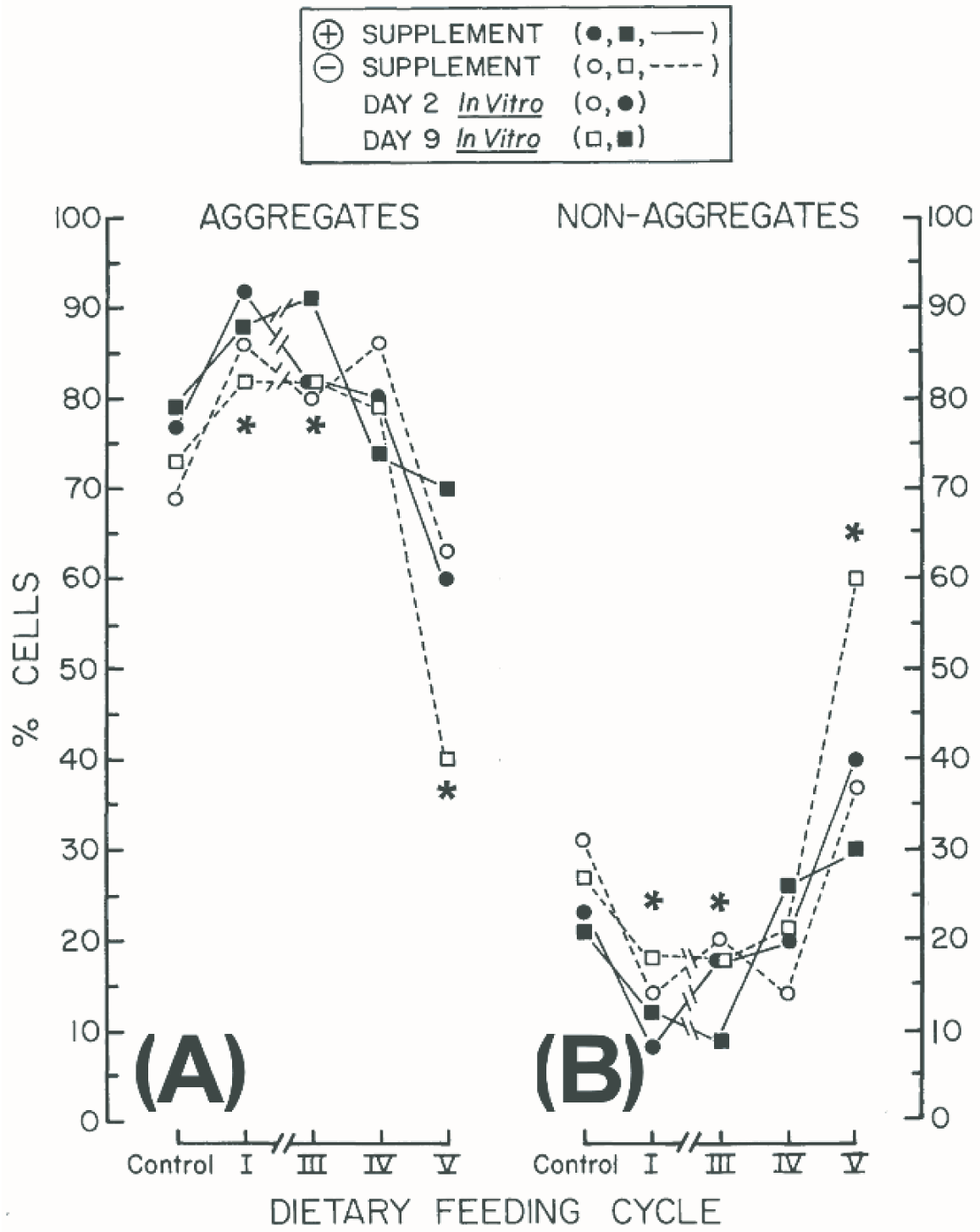
Aggregation properties of hepatocytes in primary cultures derived from rats exposed to a multi-cycle AAF feeding regimen. The percentages (%) of cells in aggregates were determined from the average numbers of hepatocytes in four boxed areas, where each box = 200 x 200 µm ([cells/40,000 µm^2^] ÷ 4), where theoretical occupation is 1 cell/400 µm^2^ at the center of each dish. The percentages (%) of non-aggregated cells in were determined by subtracting % aggregates from 100. Control results (AAF^-^S^-^ or AAF^-^S^+^) from cycles I, III, IV and V were combined and averaged (N = [2 rats] x [4 cycles] x [3 dishes/point] = 24); experimental results for each point, N = [3 rats] x [3 dishes/point] = 9). Standard error bars are not shown for points where \*\*\**P* <.005 and \**P* <0.03 (cycle feeding vs. control). See phase and light-level photomicrographs in Supplementary Figures 1 and 2.

Starting with control AAF^-^ cultures, bell-shaped curves were observed in the levels of aggregation across all feeding cycles (Figure 5A); curves of non-aggregated cells showed inverse relationships (Figure 5B). No differences were observed in the aggregation behavior of AAF^-^S^-/+^ and AAF^+^S^-/+^ hepatocytes across all feeding cycles with respect to time after plating (N = 4) or supplement addition (N = 2) (Figure 5A). In contrast, compared to the levels of aggregation in control AAF^-^S^-/+^ and cycle IV AAF^+^S^-/+^ cells (74.3 ± 4.2% and 79.3 ± 5.7%, respectively [NS]), aggregation levels in AAF^+^S^-/+^ cultures peaked during cycle I and III (86.5 ± 3.8% and 83 ± 5.4% [*P\** < 0.005 and *P\** < 0.008], respectively) and then fell significantly to a nadir during cycle V (58 ± 12.7% [*P\** < 0.003]).

### Cytotoxic and Cytocidal Effects of AAF on Cultures of Primary Hepatocytes Isolated from Adult Male Rats Exposed to a Multi-Cycle AAF Feeding Regimen

Hepatocyte DNA synthesis rates and attachment were used to quantify the cytotoxicity and lethality of extracellular AAF, respectively. Figure 6 shows the results of exposure to 0.2 mM AAF on DNA synthesis rates (Figure 6A) and attached cells (Figure 6B) in cultures of AAF^-^ or cycle I, III, IV and V AAF^+^ hepatocytes, between 1-2 (top) and 8-9 days (bottom) post-plating.

**Figure 6.**
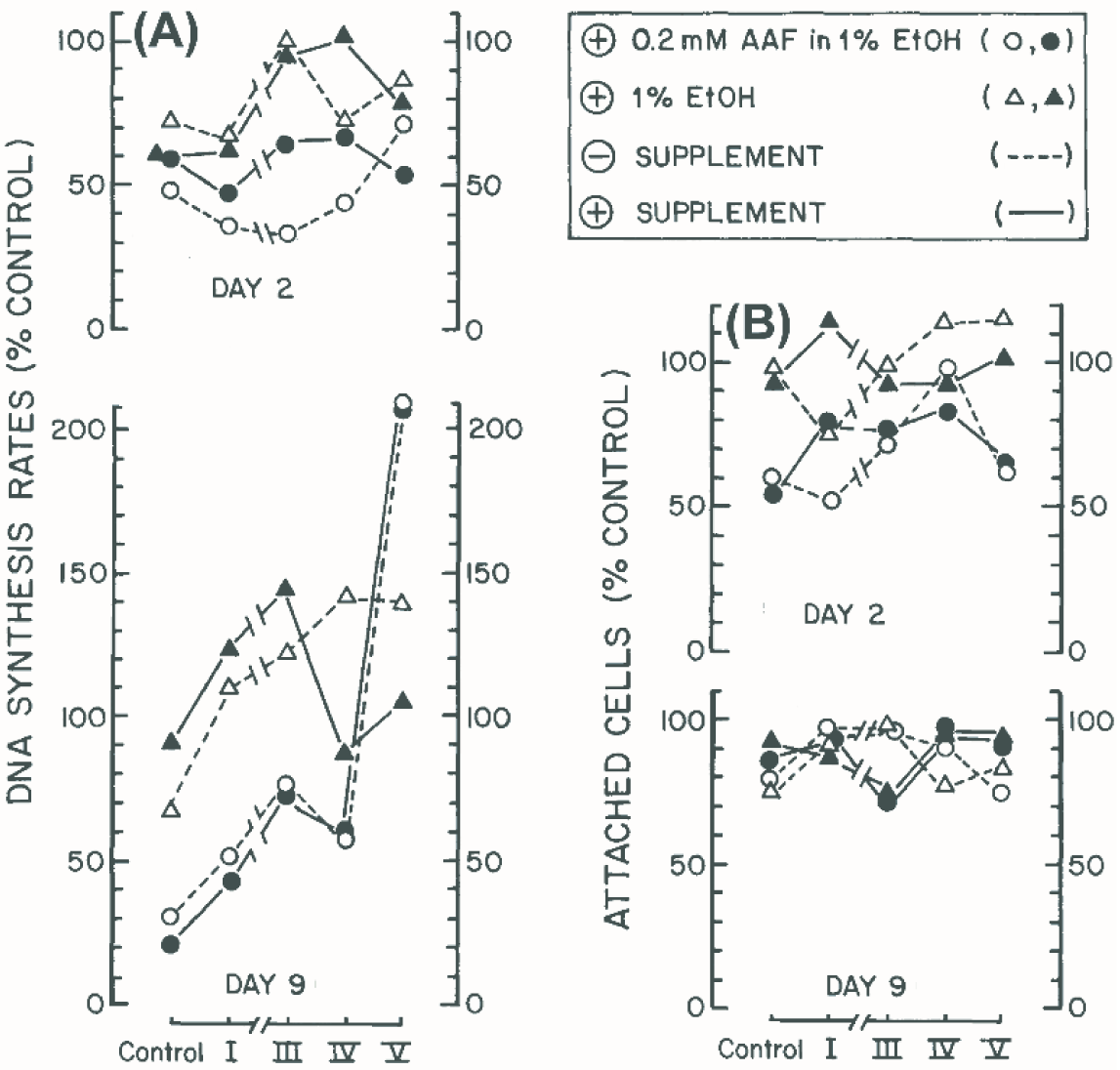
Toxicity-challen<u>ge</u> responses of hepatocytes in primary cultures derived from rats exposed to a multi-cycle AAF feeding regimen. Freshly isolated AAF^-^ and AAF^+^ hepatocytes from cycles I, III, IV and V were isolated and plated as described in Material and Methods. DNA synthesis rates ([^3^H]-TdR cpm/10^6^ cells/24 h) or attachment (cell number/dish) were measured on day 2 or day 9 following 24 h *in vitro* exposure to fresh plating media supplemented (see inset box [upper right] for symbols) with either 0.2 mM AAF in1% EtOH (○, ●) or 1% EtOH (□, □), and supplement (---) or diluent ([serum-free ornithine-supplemented media]). Control levels (x-axes) used as "100%" levels (y-axes) were taken from results obtained in Figure 4 with AAF^-^ cultures to which nothing was added: DNA synthesis rates (Figure 4B), 2000 on day 2 (S^-/+^), to 15000 on day 9 (S^-^) and 23,000 on day 9 (S^+^); and, attached cells (Figure 4C), 1.1 x 10^5^ (S^-^) and 2.2 x 10^5^ (S^+^) on day 2, to 2.2 x 10^5^ (S^-^) and 5.7 x 10^5^ (S^+^) on day 9. Control results (AAF^-^S^-^ and AAF^-^S^+^ [□- - - - □ and ▪- - -▪, respectively]) from cycles I, III, IV and V were combined and averaged (N = [2 rats] x [4 cycles] x [3 dishes/point] = 24); and plotted at the left but together with each set of experimental curves (where each point, N = [3 rats] x [3 dishes/point] = 9).

Control DNA synthesis rates (S^-/+^) were inhibited by AAF and by the AAF vehicle (1% EtOH) an average of 40% on day 2; in contrast, AAF exerted significantly more inhibition (75%) than EtOH (20%) on day 9 (Figure 6A). Inhibition patterns observed in day 2 AAF^+^ cultures from feeding cycles I-V (top) only marginally differed from controls: AAF or EtOH inhibited DNA synthesis rates an average of 60-40% or 35-20%, respectively (S^-/+^). Day 9 cultures showed completely different patterns of inhibition: AAF^+^ hepatocytes became less sensitive to the cytotoxicity of either chemical in proportion to the numbers of feeding cycles *in vivo*. Thus, in AAF-treated cultures (bottom), inhibition (S-/+) fell from control levels of 75% to 55, 35 and 40%, and then to 210% (cycles I, III and IV, or V, respectively), while inhibition rates in EtOH-treated cultures rose to ∼115-130% (S-/+), and then either leveled off at ∼140% (S-) or oscillated (S+) slightly from ∼15 to 105% (cycles I, III or IV to V, respectively).

Cell attachment (Figure 6B) in day 2 (top) or day 9 (bottom) control cultures (S^-/+^) was affected negligibly by EtOH treatment (0 or 20%, respectively), whereas AAF treatment (S^-/+^) inhibited attachment 50% on day 2, but only 20% on day 9. In cycle I, III, IV and V cultures, EtOH had little effect on day 2; but attachment fell 20-40% in AAF treated cultures (S^-/+^). On day 9, both agents similarly inhibited attachment 10-30% in cycle I-V cultures (S^-/+^).

### Pathophysiological Changes in the Premalignant Livers of Adult Rats Fed a Discontinuous 5-cycle Dietary Regimen Containing AAF

The carcinogenic AAF regimen used here differed slightly from the original model (0.06% AAF for 4 cycles, where 1 cycle = 3 week plus AAF, 1 week minus AAF). Yet a similar pattern of pathological changes was observed (reduced food consumption, induced weight loss), as evident by the early appearance of AFP^+^ oval cells and the concomitant elevation of serum AFP, followed by the appearance of hepatocellular AFP^-^ foci, AFP^-^ nodules and well-defined AFP^+^ hepatocellular carcinomas HCCs (_).

Such typical histological changes have been documented extensively (_-_). In contrast to cell culture findings of increased cell proliferation, hepatocyte proliferation is blocked during cycle I, and scattered basophilic foci of atypical AFP^-^ hepatocytes are observed. Periportal oval cells proliferate during cycle III and spread throughout AFP^-^ hepatocellular cords; many are AFP^+^. Concurrently, through cycles III-V, plasma AFP levels rise exponentially to peaks of 5 µg/mL. Hyperplastic AFP^-^ nodules (0.1-0.5 cm) emerge during cycle IV frequently in the left and median lobes; nodule numbers and sizes crest and double by cycles IV and V, respectively. Intralobular AFP- and AFP+ duct-like structures form between cycles IV-V, after which nodules regress almost entirely. When AAF^-^ diets are restored to 21-week-old AAF^+^ rats, AFP levels decline to baseline (50 ng/mL) by week 40. HCCs usually appear between 48-52 weeks; 50-60% of the primary tumors are AFP^+^. Cells from primary tumors have been cloned; many cell lines are transplantable and retain morphologies of AFP^+^ HCCs.

## DISCUSSION

Investigations with primary cultures of normal adult rat hepatocytes have led to the following working hypotheses (Koch *et al*., 2018a,b): AAF diffuses by 1^st^-order kinetics throughout bilayer lipid membranes where it encounters Cyp1A2 molecules associated with *two different compartments*. Lineweaver-Burk analyses (Koch *et al*., 2018a) of secreted AAF metabolites, and studies with isolated hepatocyte nuclei (Koch *et al*., 2018b), suggest that Cyp1A2 metabolism systems exist both in the nucleus (designated System I) and the ER (designated System II). Four principal constants defined these systems: an high-affinity K_m[APPARENT]_ (1.64 x 10^-7^ M) and a low-velocity V_MAX[APPARENT]_ (0.1 nmols/10^6^ cells/day); plus a low-affinity K_m[APPARENT]_ (3.25 x 10^-5^ M) and an high-velocity V_MAX[APPARENT]_ (1000 nmols/10^6^ cells/day), respectively.^1,2^ These constants have not yet been measured over altered hepatocellular growth states *in vitro* or *in vivo*.

Reactive metabolites of AAF covalently bind to nuclear and cytoplasmic macromolecular binding sites, as revealed by autoradiography (see Figure 6, Koch *et al*., 2018a), gel electrophoresis and studies with isolated hepatocyte nuclei (see Figures 4-6, and Table 1, Koch *et al*., 2018b). Four pharmacokinetic constants have been observed that describe these events, as revealed by Scatchard analyses (see Figure 4, Koch *et al*., 2018b). These sites are attributed to genomic DNA (Site I), defined by an high-affinity K_D[APPARENT]_ (2-4 x 10^-6^ M) and low-capacity B_MAX[APPARENT]_ (6 pmol/10^6^ cells/24 h); and, to cytoplasmic (plus secreted) proteins (Site II) ranging in M_r_ 13.7 kDa – 68 kDa, defined by a low-affinity K_D[APPARENT]_ (1.5 x 10^-3^ M) and an high-capacity B_MAX[APPARENT]_ (350 pmol/10^6^ cells/24 h ), respectively. The levels of all four binding constants vary with respect to growth cycle and culture age *in vitro*. Each parameter displays a distinct pattern from early log phase (day 2 post-plating) through stationary phase (day 12 post-plating). Thus, after 2 d of invariant change, the low-capacity B_MAX[APPARENT]_ and high-affinity K_D[APPARENT]_ levels of Site I significantly *fall* from 60 to 2 picomoles/10^6^ cells/24 h (↓ 30-fold) and *rise* from 3 x 10^-5^ M to 5 x 10^-7^ M (↑ 50-fold), respectively. In contrast, the high-capacity B_MAX[APPARENT]_ and low-affinity K_D[APPARENT]_ levels of Site II modestly *rise* from 200 to 420 picomoles/10^6^ cells/24 h (↑ 2.1-fold), and *fall* from 1 x 10^-3^ to 3 x 10^-3^ M (↓ 3-fold), respectively (see Figure 1S, Koch *et al*., 2018b).^3^

In summary, two sets of four pharmacokinetic constants (a total of eight) describe Cyp1A2 processing of AAF and the binding of electrophilic metabolites of AAF in primary cultures of living rat hepatocytes isolated from animals fed normal chow diets (Koch *et al*., 2018a,b).

Experiments in this paper addressed the question, ‘Do the *in vitro* measurements thus far reflect hepatic tissue status during AAF-induced hepatocarcinogenesis’? These issues stem from investigations using primary (1^○^) rat hepatocytes and high specific activity [^3^H]-AAF that suggest hepatocytes contain 2 different subcellular compartments of Cyp1A2 that enzymatically convert AAF into reactive compounds that preferentially bind covalently to genomic DNA (in the nucleus) at low extracellular AAF (6 μM [B_MAX_∼6 pmol/10^6^ cells/24 h]), and to large numbers of proteins (in the cytoplasm) with a K_D_ ∼ 1.5 x 10^-3^ M (B_MAX_ ∼350 pmol/10^6^ cells/24 h).

As concluded above (see Impact Statement), striking similarities are observed between both Site I and Site II B_MAX_ and K_D_ expression curves during *in vitro* and *in vivo* premalignant growth transitions. These new findings elegantly fit earlier ones from the laboratory of M.C.

Poirier (Huitfeldt H.S et al. 1990), based upon direct evidence from two-color immunofluorescence identification of hepatocyte DNA and AAF-adducts, that hepatocytes growing during carcinogen exposure manifest fewer intranuclear AAF-DNA adducts. Our findings here would indicate ‘fewer and fewer’. But how these phenomena lead to malignancy remain puzzling.

## Supporting information

Supplementary Figure 1. Phase microscopy. EMT-like transitions Days 2, 4 and 18. NEED PERMISSION from Hepatology (1982).

## SUPPLEMENTARY DATA

Supplementary data are available at *Toxicological Sciences* online.

## ACKNOWLEDGMENTS

We thank Hal Skelly for technical assistance.

## FOOTNOTES

1 A third processing system is observed (AAF → 2-aminofluorene [AF]) that is unrelated to Cyp1A2. It displays the lowest but intermediate levels of K_m_ (9.6 x 10^-5^ M) and V_MAX_ (4.7 nmols/10^6^ cells/day), respectively; and is probably an arylacetamide deacetylase (Koch *et al*., 2018a).

2 Nuclear localization of System I Cyp1A2 molecules is predicted from biochemical (Koch *et al*., 2018b) and microscopic observations (Koch *et al*., 2018a), and from the presence of a C-terminal nuclear localization signal (*gmgkrrcige*) homologous to a similar region of cytochrome P-450 4 (Quattrochi LC, *et al*., 1986), but absent from all other reported cytochrome P-450s (see Figure 3S, Koch *et al*., 2018a).

3 Temporal and quantitative patterns observed in the four macromolecular binding site constants, including properties of hepatocytes measured in Figures 3 – 6, are probably regulated by mechanisms other than those that guide growth state-dependent expression of hepatocyte differentiation. For instance, the patterns observed here differ from the growth state-dependent ‘U shape’ patterns of differentiated hepatocyte functions (*e.g.*, L-type (I) pyruvate kinase, glutathione-S-transferase-B, alcohol dehydrogenase and gluconeogenesis-from-lactate) reported previously over 0-12 day intervals *in vitro*.

---

**Supplementary Figure 1.**
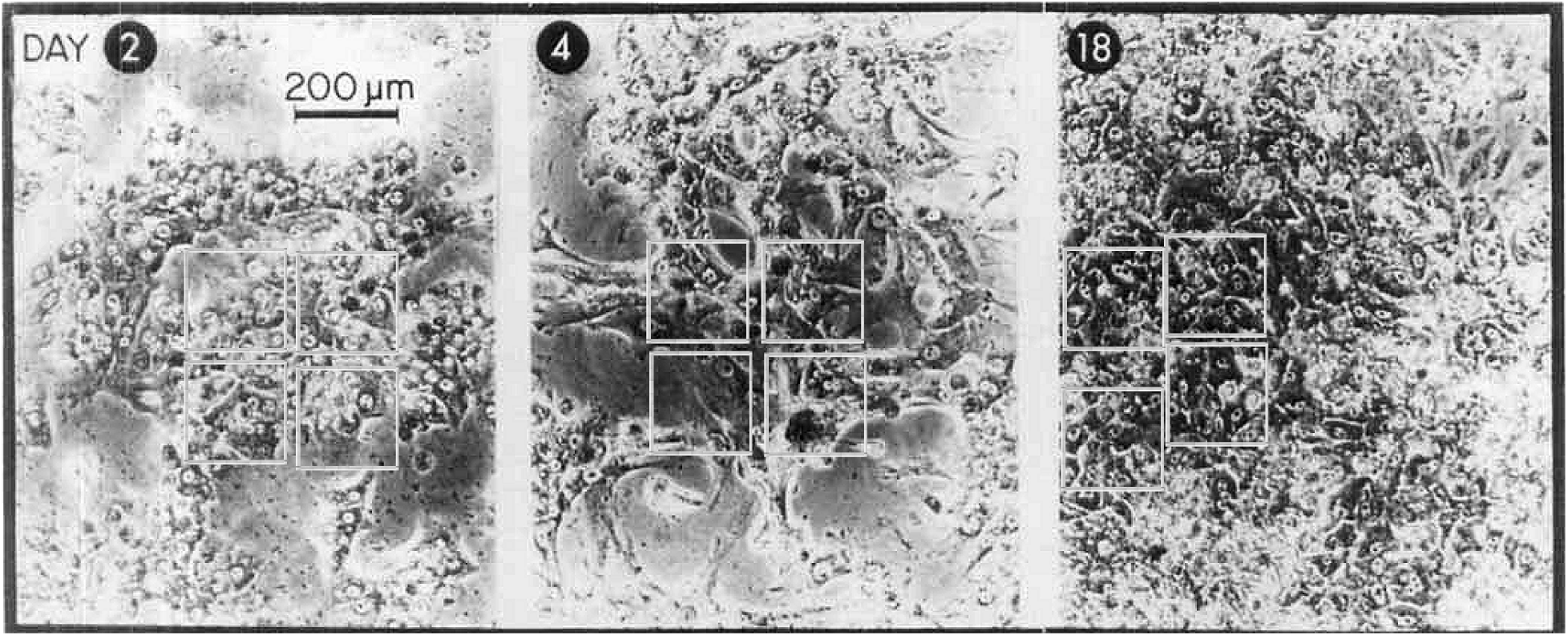
Phase microscopy. EMT-like transitions Days 2, 4 and 18. NEED PERMISSION from Hepatology (1982 paper).

