## Supplementary Figure 1. Phase microscopy. EMT-like transitions Days 2, 4 and 18. NEED PERMISSION from Hepatology (1982). for "N-Acetyl-2-Aminofluorene (AAF): *In vivo* temporal expression patterns of growth cycle-dependent macromolecular binding constants K_D[APPARENT]_ and B_MAX[APPARENT]_ are revealed by primary cultures of premalignant hepatocytes derived from a multi-cyclic hepatocarcinogenic feeding regimen"

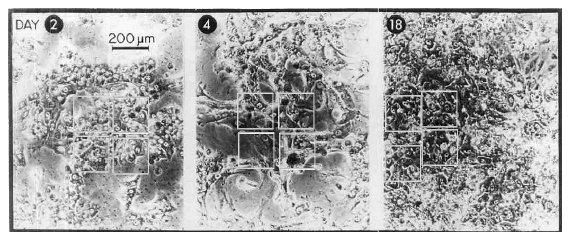
NEED PERMISSION from Hepatology (1982 paper).
